# The stability of thought: using experience sampling and brain imaging to determine the contextually bound nature of human cognition

**DOI:** 10.64898/2026.04.09.717548

**Authors:** Louis Chitiz, Samyogita Hardikar, Ian Goodall-Halliwell, Raven Wallace, Bridget Mullholland, Samuel Ketcheson, Bronte McKeown, Michael Milham, Ting Xu, Daniel Margulies, Nerissa Ho, Theo Karapanagiotidis, Giulia Poerio, Robert Leech, Elizabeth Jefferies, Jonathan Smallwood

## Abstract

Human behavior is highly flexible, allowing efficient performance across a wide range of task contexts. A distributed set of frontal and parietal regions, commonly termed the multiple-demand network (MDN), is consistently engaged during diverse cognitively demanding tasks and is thought to support this flexibility. However, it remains unclear how patterns of MDN engagement relate to the qualitative features of ongoing cognition experienced during task performance. To address this issue, we examined the reliability of self-reported experiential features sampled during performance of a broad range of tasks. Across tasks, we found little evidence that particular patterns of thought were intrinsically more reliable than others, nor that individual tasks were associated with stable, characteristic thought profiles. Instead, the reliability of specific experiential features varied systematically across task contexts, with the same patterns showing high stability in some tasks and low stability in others. We next asked whether stable patterns of thought were associated with distinct neural signatures. We found that patterns of brain activity resembling the MDN tended to be present for tasks in which deliberate task focus was high, and when distraction was lower, adding to an emerging body of research suggesting that coordinated activity within frontal and parietal regions helps to establish a stable goal-focused mode of thoughts and actions.

## INTRODUCTION

Human behavior spans a wide range of activities that differ in their cognitive demands^1–3^. Some activities require our undivided focus on an external task^4^, others rely on internally generated cognition, such as imagination or mental simulation of distant times and places^5^. Even during task performance, cognition is often unstable, with attention drifting towards unrelated concerns^6,7^. While psychology and neuroscience have identified neural systems associated with particular cognitive functions, we have a more limited understanding of how different modes of thought are organized across everyday behavior, and how the *dynamics* of these modes—their stability or variability over time—shift with situational demands.

Many forms of adaptive behavior depend on the sustained maintenance of a particular cognitive mode, such as holding task goals online, prioritizing task-relevant information, or resisting distraction over time. In other situations, less constrained or more variable patterns of thought may be beneficial. For instance, sustained attention tasks require stable cognitive states^8^, whereas creative tasks can benefit from a broader range of mental content^9^. Understanding when and how cognition is stabilized—or remains flexible—is therefore central to understanding how the brain supports effective behavior across diverse contexts^10^.

Establishing normative patterns of thought dynamics also has clinical importance^10^. Many psychiatric symptoms involve disruptions in how thoughts evolve over time. In some cases, thoughts may shift too little when variability would be adaptive (e.g., rumination, obsession, circumscribed interests, hyperfixation)^11,12^. In others, thoughts may change too rapidly when stability is required (e.g., concentration difficulties, racing thoughts)^13,14^. Characterizing how thought typically fluctuates across contexts may therefore provide a useful benchmark for identifying maladaptive deviations.

Another central unresolved issue is whether patterns of ongoing thought are primarily context-bound or reflect domain-general modes of cognition that recur across tasks. Some experiential features appear tightly linked to specific activities, such as enhanced sensory processing during movie watching^15^, while others, such as sustained task focus, may support performance across many different cognitive contexts^16^. Neuroimaging evidence points to a candidate domain-general system: a set of frontal and parietal regions that are consistently recruited across diverse tasks despite large differences in their surface features. This frontoparietal network, commonly referred to as the multiple-demand system^4^, is widely thought to support general aspects of cognitive control. However, the contribution of this system to the organization of ongoing experience remains unclear. Although the multiple-demand system is often discussed in terms of task difficulty or flexible control, an alternative possibility is that coordinated activity within frontal and parietal cortex supports the stabilization of task-appropriate modes of thought over time. From this perspective, multiple-demand engagement may be linked not only to what task is being performed, but to how consistently a particular cognitive state is sustained during performance.

The goal of the current study was to determine how patterns of ongoing thought are organized across different task contexts, and how stable these patterns are within and across tasks. We analyzed a previously published dataset^17^ in which 200 participants provided repeated experience-sampling reports while performing 14 distinct tasks. This design allowed us to quantify the reliability of different experiential features over time and to ask whether stability is a property of particular thought patterns, particular task contexts, or their interaction. Finally, we examined whether tasks associated with more stable patterns of thought were characterized by distinct profiles of brain activity, allowing us to link experiential stability to large-scale neural organization (see Fig. 1 for a schematic of our analytic pipeline).

**Figure 1.**
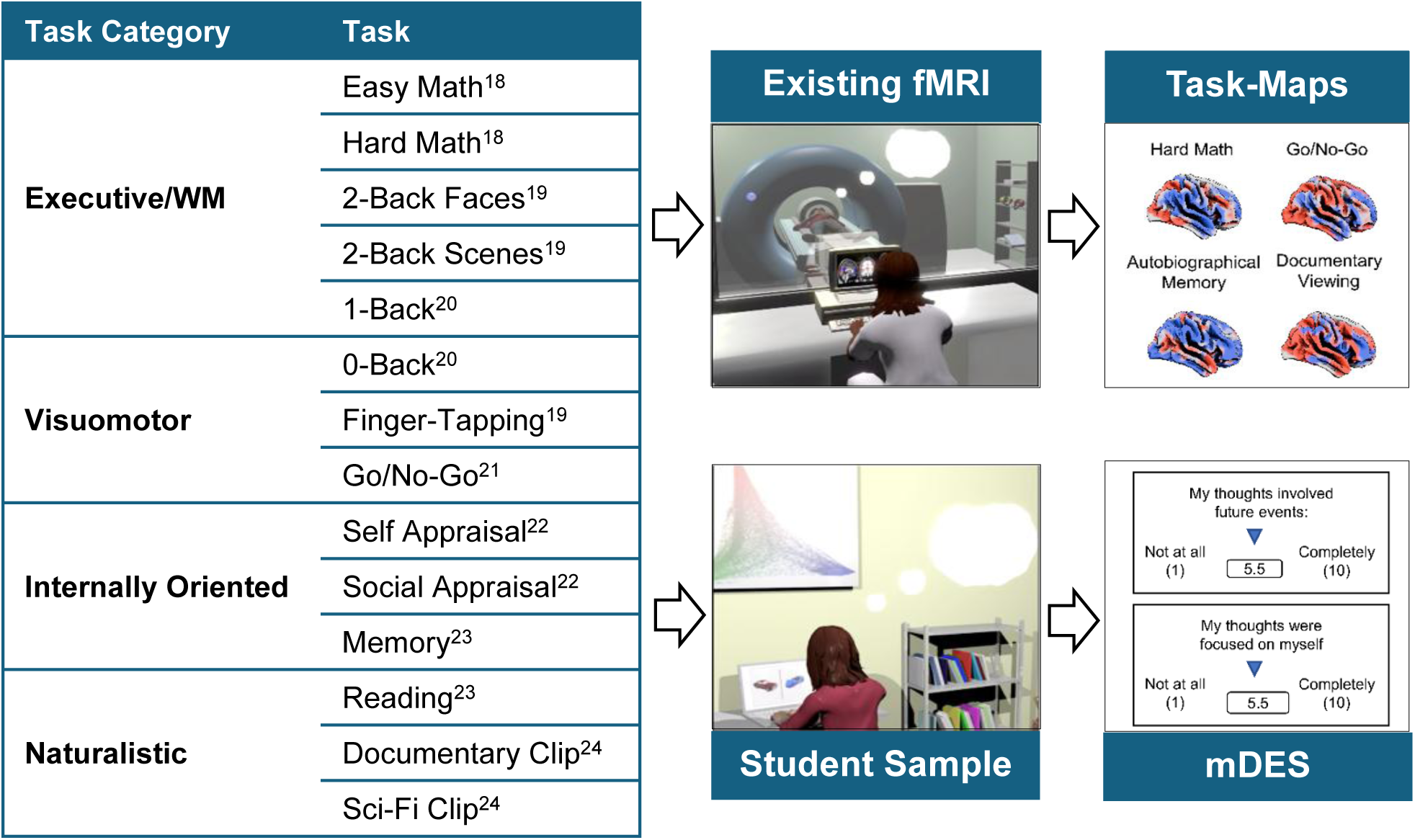
Overview of data collection. *(Left)* We selected 14 tasks to capture states linked to a range of cognitive domains. *(Right)* To represent the brain activity and internal experience associated with each task, we obtained unthresholded task maps from existing fMRI data and administered multidimensional experience-sampling (mDES) thought probes while participants completed each task.

## RESULTS

### Generating a ‘thought-space’ and ‘brain-space’

Whole-brain activation and ongoing thought each comprise a high number of dimensions making our research questions computationally difficult to investigate. To make our analyses more feasible, we used dimensionality reduction to represent each measure in lower-dimensional space: a ‘thought-space’ for participants’ mDES responses and a ‘brain-space’ for the brain activity for each task^1,2,15,17,18,25–31^. These spaces allow for a clearer representation of how different contexts compare psychologically and neurologically. For example, tasks closer together in the ‘thought-space’ tend to evoke more similar descriptions of internal experience, while tasks closer together in the ‘brain-space’ engender more similar patterns of whole-brain activity.

Per our previous work (see ^32^ for a review; ^1,2,15,25,29^), we generated our ‘thought-space’ by applying Principal Components Analysis (PCA) to the full set of thought probes across our sample. Based on examination of the Scree plot and parallel analysis we extracted 4 components, accounting for ∼49% of total variance (See Fig. 2). After applying Varimax rotation, we labelled the components based on their loading structure. The four components were: 1) *Episodic Knowledge*, reflecting reliance on past events and stored knowledge; 2) *Intrusive Distraction*, characterized by strong loadings on intrusive and distracting features; 3) *Deliberate Task-Focus*, defined by deliberate and task-relevant features; and 4) *Sensory Engagement*, reflecting imagery- and sound-based features (Fig. 2). These components each have precedent in the mDES literature (e.g., ^1,2,15,28,29^).

**Figure 2.**
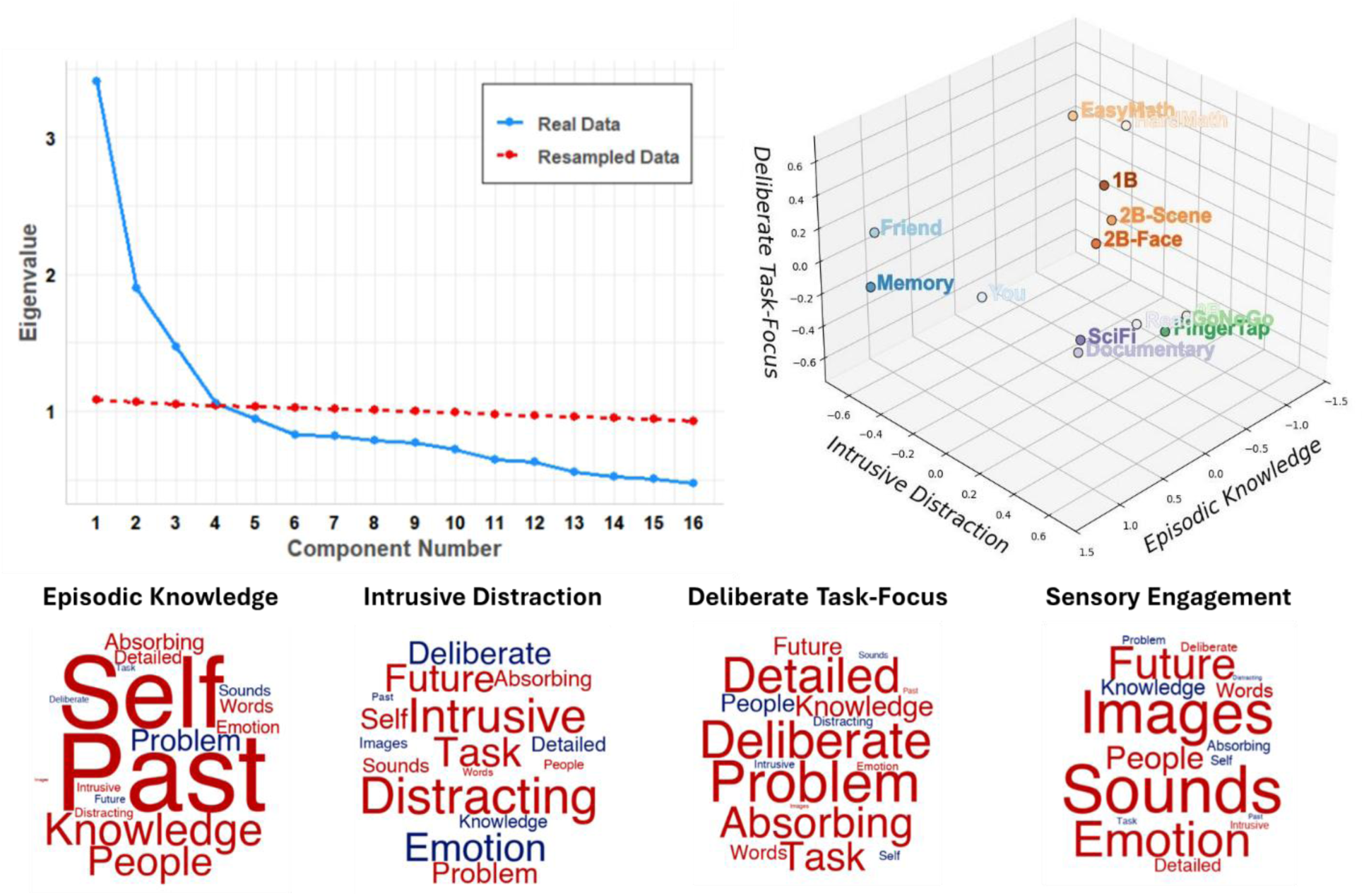
Component extraction and distribution of tasks in the ‘thought-space’. To represent participants’ reported thoughts in fewer dimensions, we decomposed the set of mDES probes using PCA. *(Top Left)* Based on examination of the Scree plot and parallel analysis we extracted 4 components. *(Top Right)* These components comprise the dimensions of a continuous ‘thought-space’ where we can compare reported thought across contexts in a data-driven manner. This 3D scatterplot depicts the average scores for each task on the first 3 Varimax-rotated components, “*2B-Face/2B-Scene*” = 2-back task with faces/scenes, “*0B/1B*” = 0-back/1-back task, “*EasyMath/HardMath*” = easy and hard math tasks, *“Read”* = reading task, “*Memory*” = memory task, “*Friend*” = social appraisal task, “*You*” = self-appraisal task, “*FingerTap*” = finger-tapping task, “*GoNoGo*” = go/no-go task, “*Documentary/SciFi*” = movie-viewing of documentary/sci-fi clips. *(Bottom)* We named the Varimax-rotated components according to their loading structure, which we visualize using Word Clouds, where word size represents loading strength (larger = stronger loading) and color represents directionality (red = positive, blue = negative).

For sake of clearer visualization, we grouped the 14 tasks in our battery according to 4 categories based on their associated cognitive demands (see Fig. 1): 1) “*executive/WM*” tasks that recruit executive functioning and working-memory (WM), 2) “*visuomotor*” tasks, which rely on basic perceptual decision-making, 3) “*internally oriented*” tasks, which require participants to attend to their inner thoughts, and 4) “*naturalistic*” tasks designed to emulate activities participants may engage in during their daily lives. These groupings were used solely for descriptive purposes and were not used in any statistical analyses.

Figure 3 shows the distribution of the tasks in our battery on each component. Participants scored highest on *Episodic Knowledge* during *internally oriented* tasks such as a cued autobiographical memory task (“Memory task”; *mean*= 1.64, 95% *CI*[1.54, 1.73]), or while deciding if character traits applied to a close friend (“Friend task”; *mean*= 1.28, 95% *CI*[1.20, 1.37]), or oneself (“You task”; *mean*= 1.19, 95% *CI*[1.10, 1.28]). *Episodic Knowledge* scores were lowest during *visuomotor* tasks such as the response-inhibition go/no-go task (“GoNoGo”; *mean*= −0.71, 95% *CI*[−0.83, −0.60]) and the motoric decision-making finger-tapping task (“FingerTap”; *mean*= −0.58, 95% *CI*[−0.70, −0.47]). *Intrusive Distraction* was reported most frequently during *visuomotor* tasks such as when matching basic shapes from a set (“0-Back task”/“0B”; *mean*= 0.47, 95% *CI*[0.35, 0.58]) and while reading (“Read”; *mean*= 0.32, 95% *CI*[0.21, 0.44]). *Intrusive Distraction* appeared the least during the *internally oriented* Friend task (*mean*= −0.52, 95% *CI*[−0.63, −0.42]) and Memory task (*mean*= −0.35, 95% *CI*[−0.47, −0.24]).

**Figure 3.**
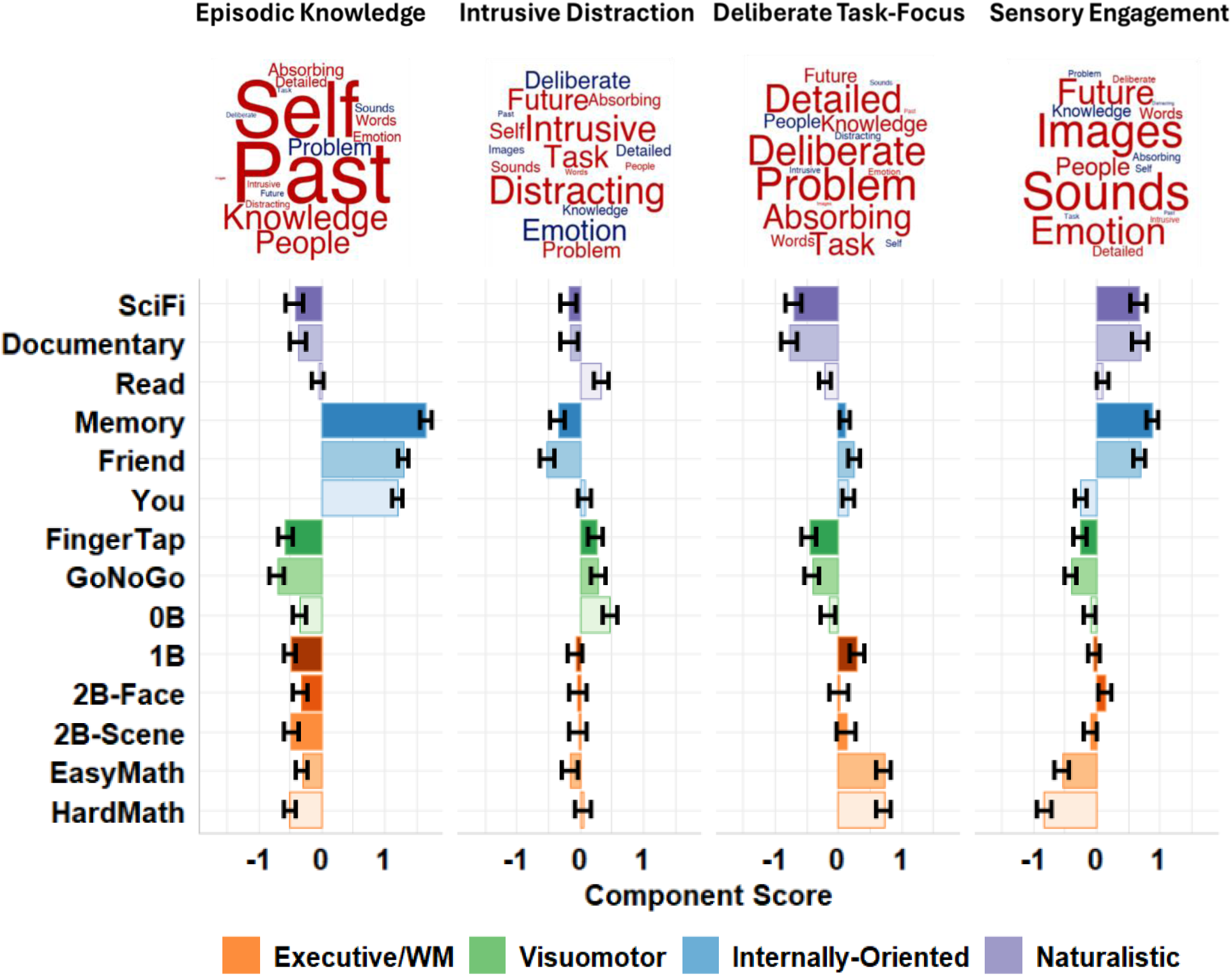
Distribution of tasks on each component. *(Top)* We named the Varimax-rotated components according to their loading structure, which we visualize using Word Clouds, word size represents loading strength (larger = stronger loading) and colour represents directionality (white = positive, black = negative). *(Bottom)* The distribution of tasks on each component based on average component score, error bars represent mean±SEM, “*2B-Face/2B-Scene*” = 2-back task with faces/scenes, “*0B/1B*” = 0-back/1-back task, “*EasyMath/HardMath*” = easy and hard math tasks, *“Read”* = reading task, “*Memory*” = memory task, “*Friend*” = social appraisal task, “ *You*” = self-appraisal task, “*FingerTap*” = finger-tapping task, “*GoNoGo*” = go/no-go task, “*Documentary/SciFi*” = movie-viewing of documentary/sci-fi clips.

*Executive/WM* tasks such as the fast mental addition during the easy math (“EasyMath”; *mean*= 0.71, 95% *CI*[0.59, 0.83]) and hard math tasks (“HardMath”; *mean*= 0.71, 95% *CI*[0.59, 0.83]) associated with the highest scores on *Deliberate Task-Focus*, while *naturalistic* passive viewing of a documentary clip (“Documentary”; *mean*= −0.77, 95% *CI*[−0.90, −0.65]) and sci-fi clip associated with the lowest scores (“SciFi”; *mean*= −0.72, 95% *CI*[−0.85, −0.59]). Finally, participants reported the highest *Sensory Engagement* during the Memory task (*mean*= 0.86, 95% *CI*[0.77, 0.96]) and the lowest during the hard math task (*mean* = −0.84, 95% *CI*[−0.95, −0.73]).

To represent the brain activation associated with each task we used group-level unthresholded maps from existing fMRI datasets (see Fig. 1 for sources). We projected them onto an existing decomposition of a larger resting-state dataset from the Human Connectome Project (HCP)^19,33^ which we use as a low-dimensional brain space. This decomposition used diffusion-embedding to identify large-scale patterns or “gradients” in whole-brain functional connectivity, in descending order of variance accounted for^33^ (see Methods). Each gradient captures a different way that brain areas can organize into anti-correlated functional networks (e.g., sensorimotor areas versus transmodal areas). These dimensions are termed “gradients” because they order brain regions along a spectrum between the two networks rather than binning them into each network categorically (e.g., the principal gradient orders regions from sensory to transmodal cortex). In our analysis, we focused on the first five cortical gradients, which capture important theoretical features in the brain’s functional organization^33^ and which correlate with patterns in reported thought^17,30^ (see Fig. 4).

**Figure 4.**
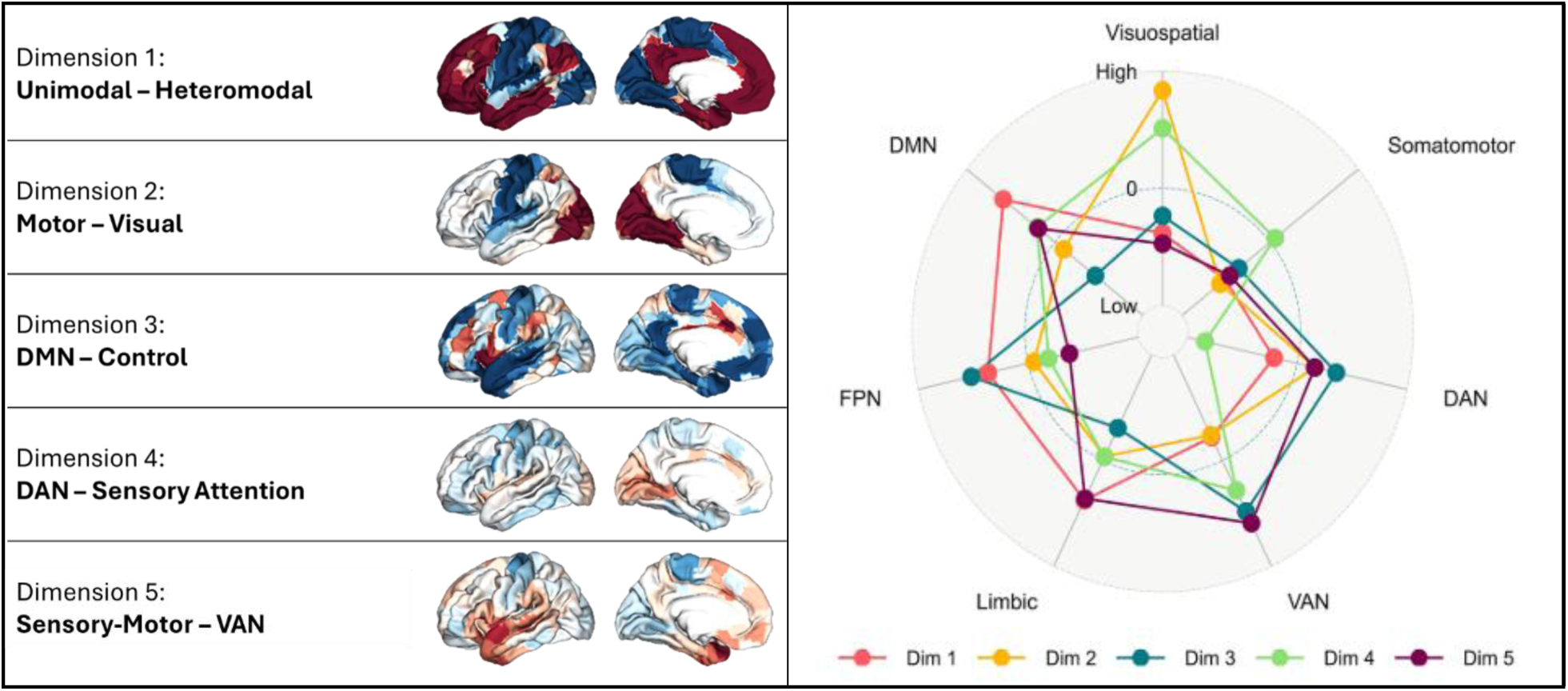
Connectivity gradients and distribution of tasks in “brain-space”. (*Left*) The five principal connectivity gradients. Regions in blue depict brain areas linked to the low end of the gradient while red regions fall to the high end. (*Right*) Adapted from ^30^; network loadings of each of the five gradients on 7 canonical resting-state networks from ^34^.

The five principal gradients are: 1) “unimodal – heteromodal”, which separates sensory from association cortex, 2) “motor **–** visual”, which splits up sensorimotor cortex into perceptual and somatomotor regions, 3) “DMN **–** control”, which divides association cortex separating the default mode network (DMN) from the frontoparietal control network (FPCN), 4) “DAN **–** sensory attention”, which separates the dorsal attention network (DAN), at one end, from visual cortex and the ventral attention network (VAN) at the other and 5) “sensory-motor **–** Limbic-VAN”, separates primary vision, at one end, from regions of auditory cortex and regions linked to language (such as the anterior temporal lobe and the left inferior frontal cortex, see Fig. 4). To project each task-map onto these 5 dimensions, we correlated the maps with each of the 5 gradients. The resulting 5 correlations per task-map represented five “coordinates” situating the functional organization of the map in ‘brain-space’ (see ^17^ for a similar approach). See Figure 5 for the distribution of tasks along each gradient.

**Figure 5.**
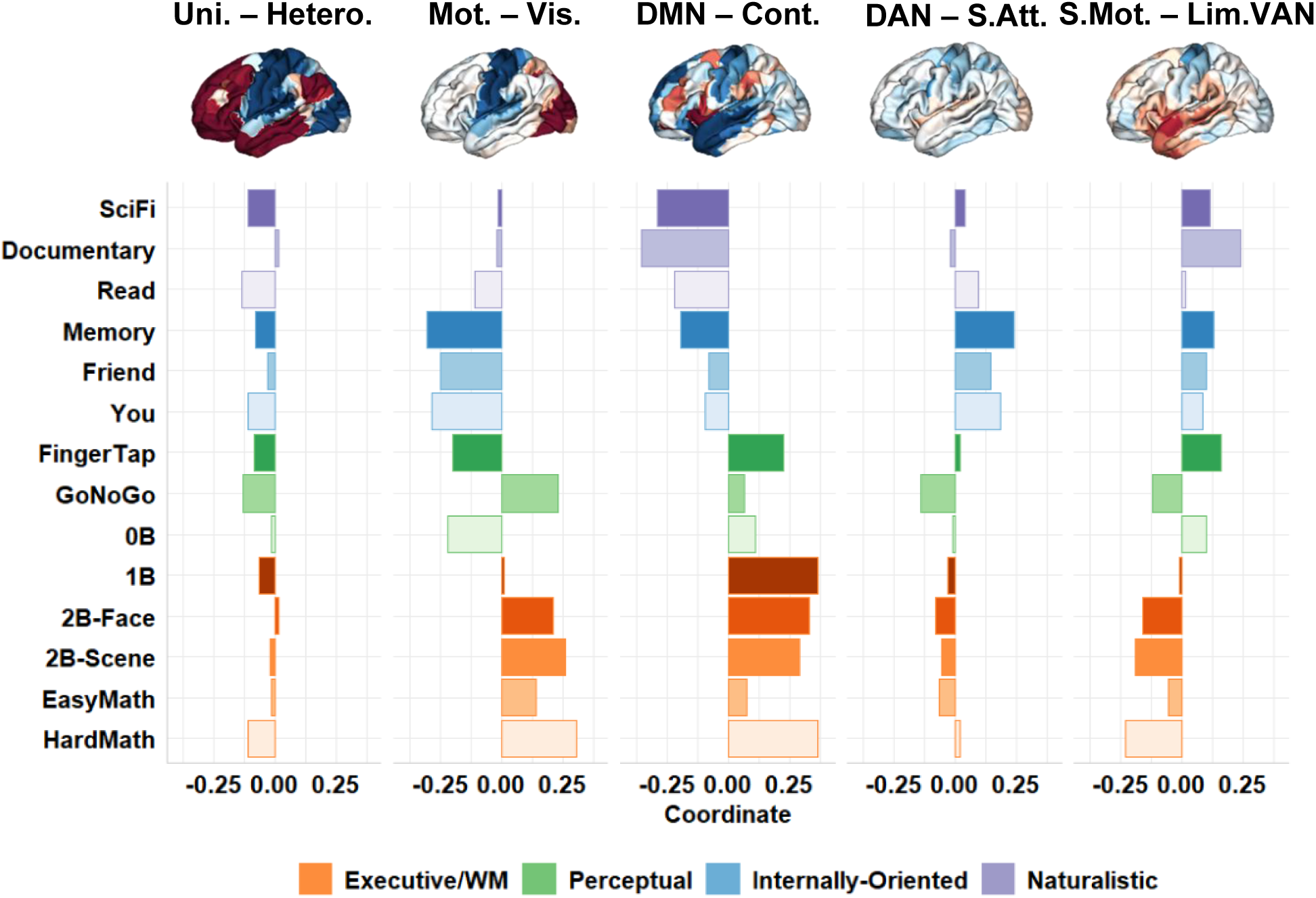
Gradient coordinates of each task. . To represent task-related whole-brain activation in fewer dimensions we projected unthresholded group-level task maps onto the first five cortical gradients generated from resting-state fMRI data from the Human Connectome Project. See here the distribut ion of the 14 tasks on each of the 5 gradients. “Uni.-Hetero.” = “Unimodal – Heteromodal”; “Mot.-Vis.” = “Motor – Visual”; “DMN-Cont.” = “DMN – Control”; “DAN-S.Att.” = “DAN – Sensory Attention”; “S.Mot.-Lim.VAN.” = “Sensory-Motor – Limbic-VAN”.

### Trait versus state influence on internal experience changes depending on task and thought content

Having reduced the number of dimensions representing each measure, our next analysis set out to examine how variance in each thought pattern could be attributed to dependencies related to subject and/or task. We quantified reliability at three levels. A set of ICCs was first estimated from intercept-only mixed-effects models to characterise the intrinsic reliability of each component when variance was partitioned only by subject or by task. These null ICCs provide reference points: within-subject ICCs index how consistently individuals experienced each component of thought across all tasks, whereas within-task ICCs index how consistently each task elicited a given component of thought across participants. We then examined stability at the level of specific task-by-subject pairs for each component. For these analyses, ICCs were estimated from mixed-effects models including task cross-factored with subject as random factors (i.e., probes were nested within task by subject combinations). In this framework, the ICC represents the expected correlation between two probes sampled from the same participant performing the same task, allowing us to quantify how strongly each task stabilised each pattern of thought (see Methods).

While certain thought patterns appeared more driven by individual differences and others more by task differences (see Supplementary Results), all 4 components demonstrated limited stability both within-subjects across tasks [*Episodic Knowledge*: *ICC*_*subject*_ = .17; *Intrusive Distraction*: *ICC*_*subject*_ = .37; *Deliberate Task-Focus*: *ICC*_*subject*_ = .33; *Sensory Engagement*: *ICC*_*subject*_ = .28] and within-tasks across subjects [*Episodic Knowledge*: *ICC*_*task*_ = .36; *Intrusive Distraction*: *ICC*_*task*_ = .04; *Deliberate Task-Focus*: *ICC*_*task*_ = .13; *Sensory Engagement*: *ICC*_*task*_ = .20] (^35^; see Table 1). In other words, reported thought was not predominantly attributable to either the individual scoring the probe or to their task.

**Table 1.** ICCs for each thought-pattern nested within-subject, within-task, or cross-factored subject:task. ICCs are presented with bootstrapped 95% confidence intervals.

| Thought-Pattern | Subject ICC | Task ICC | Subject x Task ICC |
| --- | --- | --- | --- |
| *Episodic Knowledge* | .17 [.13, .20] | .36 [.19, .55] | .66 [.64, .68] |
| *Intrusive Distraction* | .37 [.31, .42] | .04 [.00, .07] | .57 [.55, .59] |
| *Deliberate Task-Focus* | .33 [.28, .38] | .13 [.04, .20] | .68 [.66, .69] |
| *Sensory Engagement* | .28 [.23, .33] | .20 [.07, .31] | .63 [.61, .65] |

We next examined how task and subject influenced reliability in combination. That is, we assessed how participants reported different types of thought more reliably *depending on the task at hand*. Here, the ICC reflects the expected correlation between two thought reports from the same participant performing the same task. Intuitively, this combination produced higher ICC estimates than subject or task alone, with all thought patterns achieving moderate stability (See Table 1). However, not all thought patterns showed equal stability: *Deliberate Task-Focus* (*ICC*_𝑠𝑢𝑏𝑗𝑒𝑐𝑡:𝑡𝑎𝑠𝑘_ = .68), *Episodic Knowledge* (*ICC*_𝑠𝑢𝑏𝑗𝑒𝑐𝑡:𝑡𝑎𝑠𝑘_ = .66), and *Sensory Engagement* (*ICC*_𝑠𝑢𝑏𝑗𝑒𝑐𝑡:𝑡𝑎𝑠𝑘_ = .63), all had non-overlapping confidence intervals with *Intrusive Distraction* (𝐼𝐶𝐶_𝑠𝑢𝑏𝑗𝑒𝑐𝑡:𝑡𝑎𝑠𝑘_ = .57). *Intrusive Distraction*, therefore, had the lowest stability of the thought patterns identified in our study.

### How does stability in reported thought relate to task condition?

Thus far our analysis indicates that reliability in reported thought is context-dependent, varying according to interactions between trait- and state-factors characterizing different subjects and tasks. To further probe this interaction, we generated ICCs for each task and each thought component to estimate the extent to which differences between subjects accounted for variance on each thought component during each task condition (see Fig. 6).

**Figure 6.**
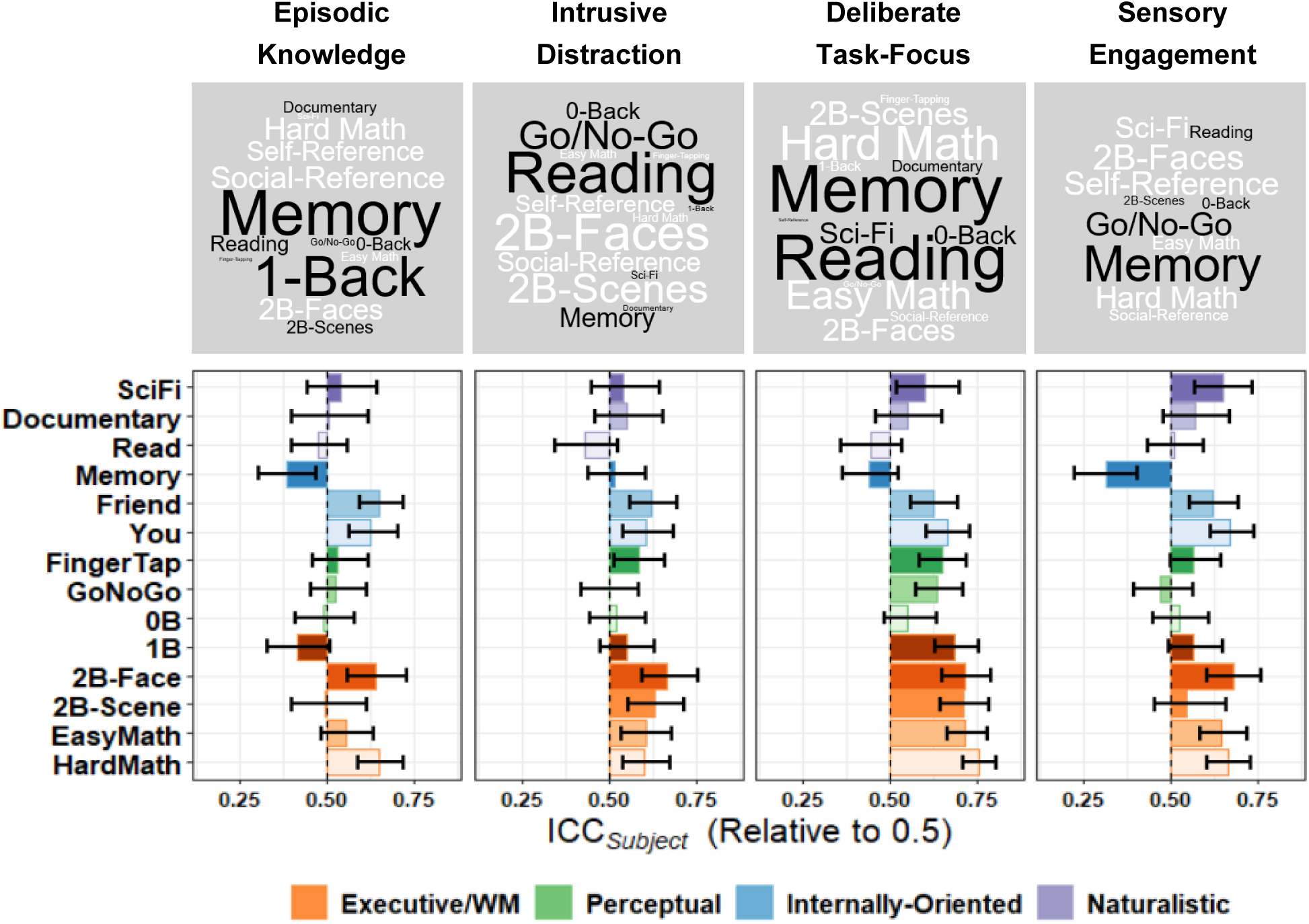
The influence of context versus subject on reported thought per task. *(Top)* Word clouds depict the distribution of mean-centred ICCs across tasks, word size indicates distance from average (larger = further from average), word color represents direction (white = above average, black = below average). *(Bottom)* While ICCs varied across tasks depending on the thought pattern in question, more challenging tasks achieved consistently higher stability across components, whereas stability on some tasks depended on the component in question. The bar plots here center around 0.50, representing the threshold beyond which subject-level dependency is apparent (i.e., subjects respond more similarly to themselves than others).

For tasks with higher ICCs, within-subject variation tended to be low while between-subject variation was high (See Fig. 7). In these cases, since participants’ probe responses tended to be consistent across probes and unique to the rest of the sample, variance in reported thought could be attributed to differences between subjects. Tasks with lower ICCs fell into two groups: 1) those where probe responses were generally unstable, such that variability was high both between and within subjects, or 2) those where responses were highly similar across participants, so that variance in reported thought was more attributable to the task than individual differences. We visualize the relationship between participant reliability and uniqueness using variation field maps^36^, which plot within- against between-individual variability for each task. As the ratio of between- to within-subject variation grows (i.e., as points move to the top left of the map), participants become more discriminable, and ICC values increase (see Fig. 7).

**Figure 7.**
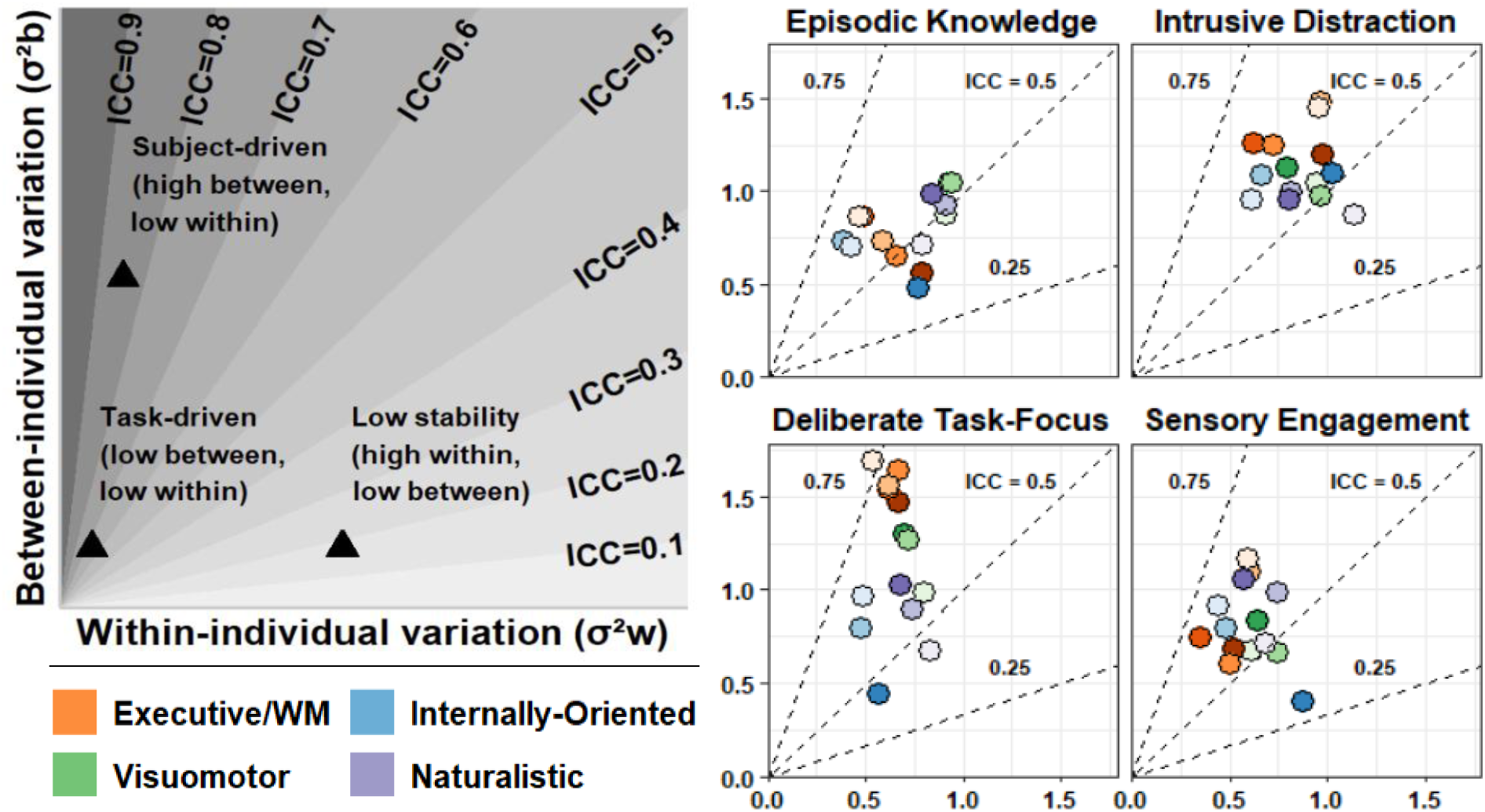
Between- versus within-individual variation across tasks. (Top Left) Adapted from ^36^; Xu’s *Theoretical Variation Field Map* visualizes how the ICC increases as individuals’ respond more consistently within themselves (i.e., lower 𝜎*_w_*^2^) and more uniquely from others (i.e., higher 𝜎*_b_*^2^), such as for the point in the top left. On the other hand, a low ICC could result from either: 1) generally unstable responses (e.g., lower right point), or 2) individuals responding too similarly to each other, becoming indistinguishable (e.g., lower left point). (*Right*) Variation field maps for each task on each thought-pattern. While passive, unengaging tasks typically demonstrated the lowest ICC values across components, field maps reveal that in some cases this was due to high similarity between subjects (e.g., the memory task) while in others it was a result of poor stabilit y overall (e.g., the reading task).

Having established that within-subject stability varied across tasks, we next examined whether this variation was related to the dominant mode of thought elicited by each task. For each component, we fit simple linear models relating task-level ICCs to the mean component score for each task. Because this analysis was necessarily conducted at the task level (14 observations), statistical power was limited. We therefore used bootstrap resampling to assess the robustness and directional consistency of each relationship rather than relying on null-hypothesis significance testing. We resampled at the participant level with replacement (𝐵 = 1000) and computed percentile-based 95% confidence intervals for each regression coefficient (see Fig. 8).

**Figure 8.**
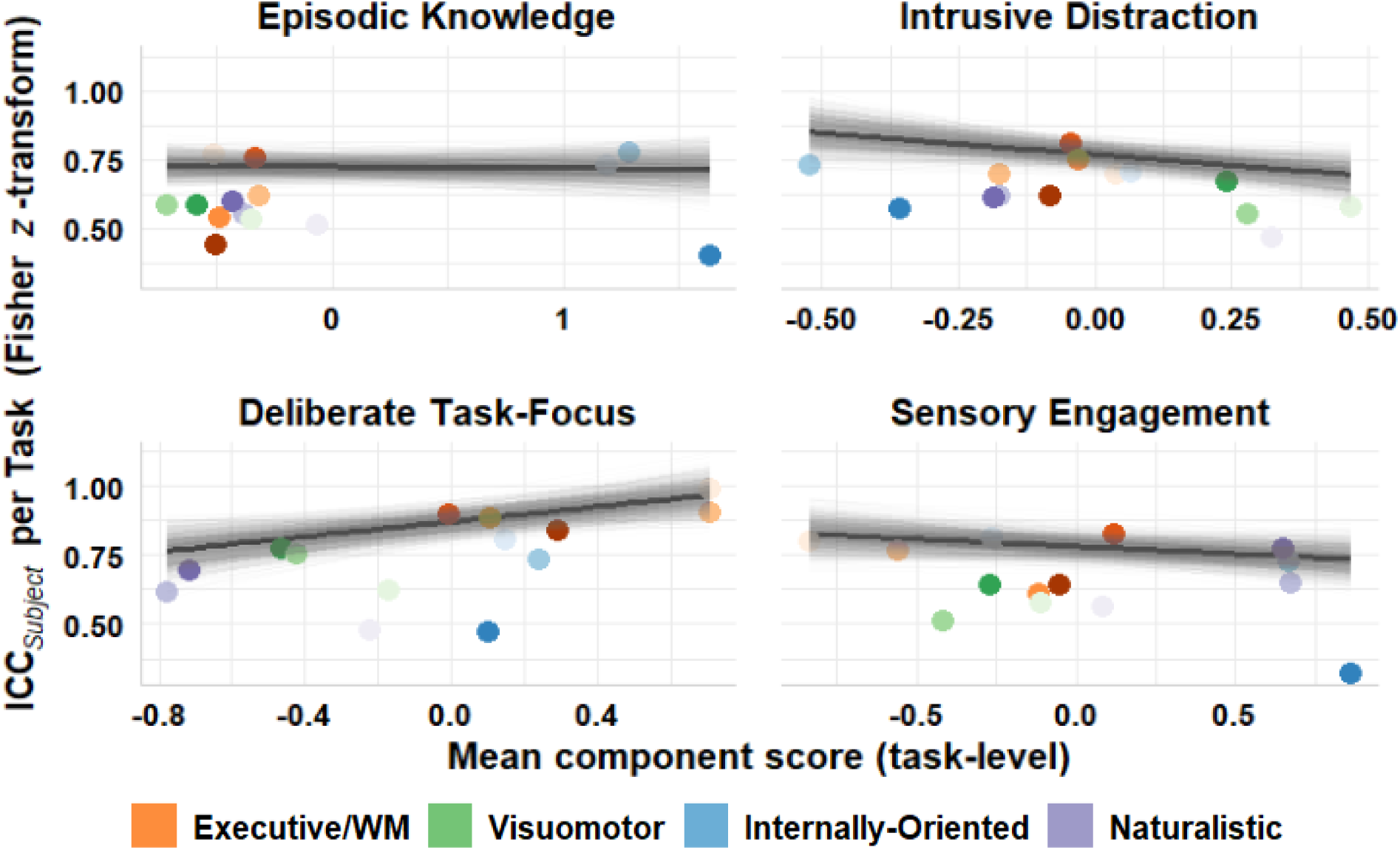
Relationship between within-subject stability and component score. To examine how within-subject stability on a given component related to its appearance, we regressed ICC per task against task-averaged component score in 1000 bootstrap resamples. Bootstrapped coefficients indicated that during tasks with high levels of task-focus, subjects’ reported thought tended to be more consistent between probes. On the other hand, tasks high in *Intrusive Distraction* or *Sensory Engagement* were associated with probe responses becoming less consistent.

Bootstrapped regression suggested that tasks with higher scores on *Deliberate Task-Focus* (i.e., with higher average levels of deliberate, externally focused thought) tended to show greater stability in probe responses. The bootstrapped median regression coefficient was positive, 𝑏 = 0.14, 95% *CI*[0.05, 0.22] . This effect was highly directionally reliable with confidence intervals excluding zero, and positive slopes emerging in 99.9% of resamples. The opposite pattern emerged for *Intrusive Distraction*, which tended to be expressed more variably across probes. Tasks with higher *Intrusive Distraction* were associated with *lower* stability in probe responses, median 𝑏 = −0.16, 95% *CI*[−0.29, −0.03]. Here too, the effect was directionally consistent across resamples with confidence intervals excluding 0, and 99% of resample estimates showing a negative slope. This was also the case for *Sensory Engagement*, though less reliably, 𝑏 = −0.05, 95% *CI*[−0.12, 0.01], suggesting probe responses tended to become less consistent in tasks evoking stronger image- and sound-based thought content. Despite confidence intervals including zero, the direction of the relationship was fairly consistent, appearing negative in 95% of resamples. There was no evidence for any reliable linear relationship between stability and component score for *Episodic Knowledge*, −0.01, 95% *CI*[−0.05, 0.04].

### What patterns of brain activity characterize tasks where thought-content is stable?

Having established how the stability of thought patterns varies across tasks, we next examined how patterns of task driven brain activity correspond to these patterns of stability on each thought pattern. Here too, we used a bootstrapping approach to mitigate power constraints from the small number of observations. We bootstrapped multiple regression models (𝐵 = 1000) resampling at the participant level with replacement. In each resample, we regressed task-level ICCs for each thought-pattern against task-level coordinates on the first five principal gradients (see Methods).

Bonferroni-corrected percentile-based bootstrapped confidence intervals (𝐶𝐼_𝑎𝑑𝑗._ = 99.75%) pointed to three reliable effects (see Fig. 9; see Table S2 for the full set of bootstrapped effects). The first set of effects suggested that subjects tended to more stably report *Deliberate Task-Focus* during tasks linked to activation of visual over somatomotor regions (*Motor* – *Visual*: median 𝑏 = 0.19, 𝐶𝐼_𝑎𝑑𝑗._[0.03, 0.34]; 100% of resamples showed positive slopes) as well as areas implicated in executive control over areas of the DMN (*DMN* – *Control*: median 𝑏 = 0.11, 𝐶𝐼_𝑎𝑑𝑗._[0.02, 0.19]; 100% of resamples showed positive slopes). Second, subjects produced more stable *Intrusive Distraction* scores during tasks linked to activation of association cortex over strictly unimodal areas (*Unimodal* – *Heteromodal*: median 𝑏 = 0.07, 𝐶𝐼_𝑎𝑑𝑗._[0.01, 0.14]; 99.9% of resamples yielded positive slopes).

**Figure 9.**
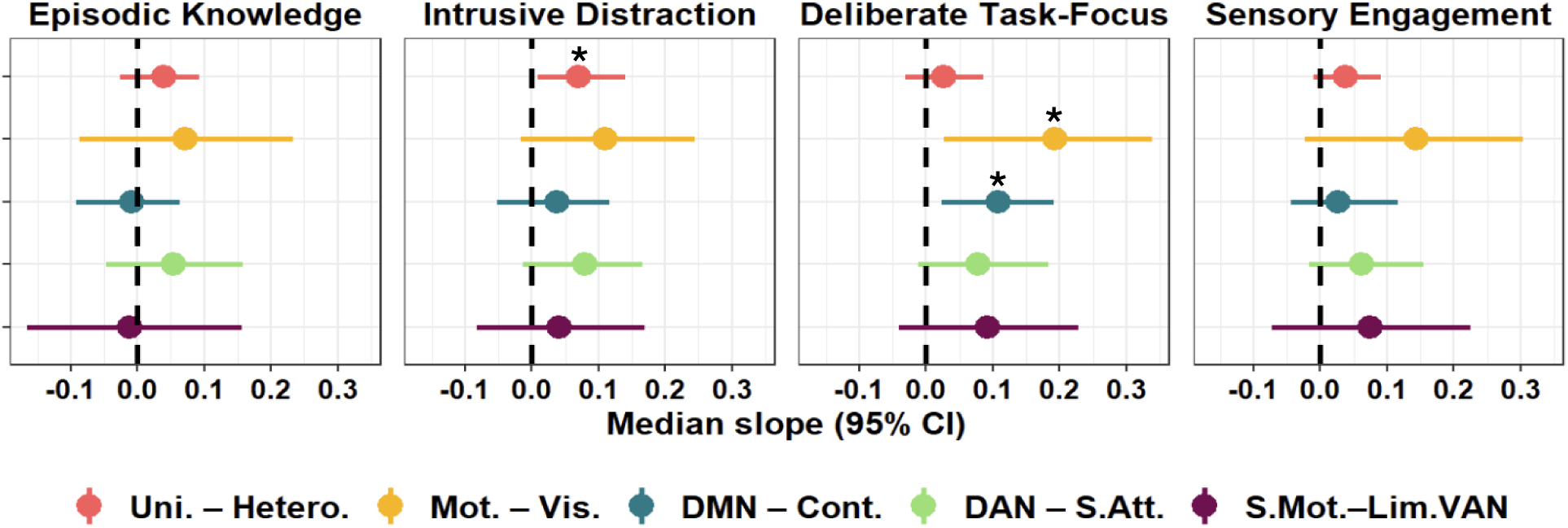
Association between stability in reported thought and cortical gradients. To examine how stability in individuals’ reported thought related to brain activation during each task, we computed bootstrapped regression coefficients representing the relationship between each gradient and stability in reported thought while holding other gradients at a constant, * = 𝐶𝐼_𝑎𝑑𝑗._ exclude 0. “Uni.-Hetero.” = “Unimodal – Heteromodal”; “Mot.-Vis.” = “Motor – Visual”; “DMN-Cont.” = “DMN – Control”; “DAN-S.Att.” = “DAN – Sensory Attention”; “S.Mot.-Lim.VAN” = “Sensory-Motor – Limbic-VAN”.

To improve interpretability for the relationships between within-subject reliability on each thought pattern and whole-brain activation in each of the tasks, we generated a set of dot-product maps for each component. For each map, we multiplied each gradient by its median bootstrapped regression coefficient on a given thought-pattern and then compressed across gradients at each parcel, generating a single ‘stability map’ (see Methods; see Fig. 10). For these stability maps, regions more strongly related to within-subject stability on the component received a stronger positive weighting and *vice versa*.

**Figure 10.**
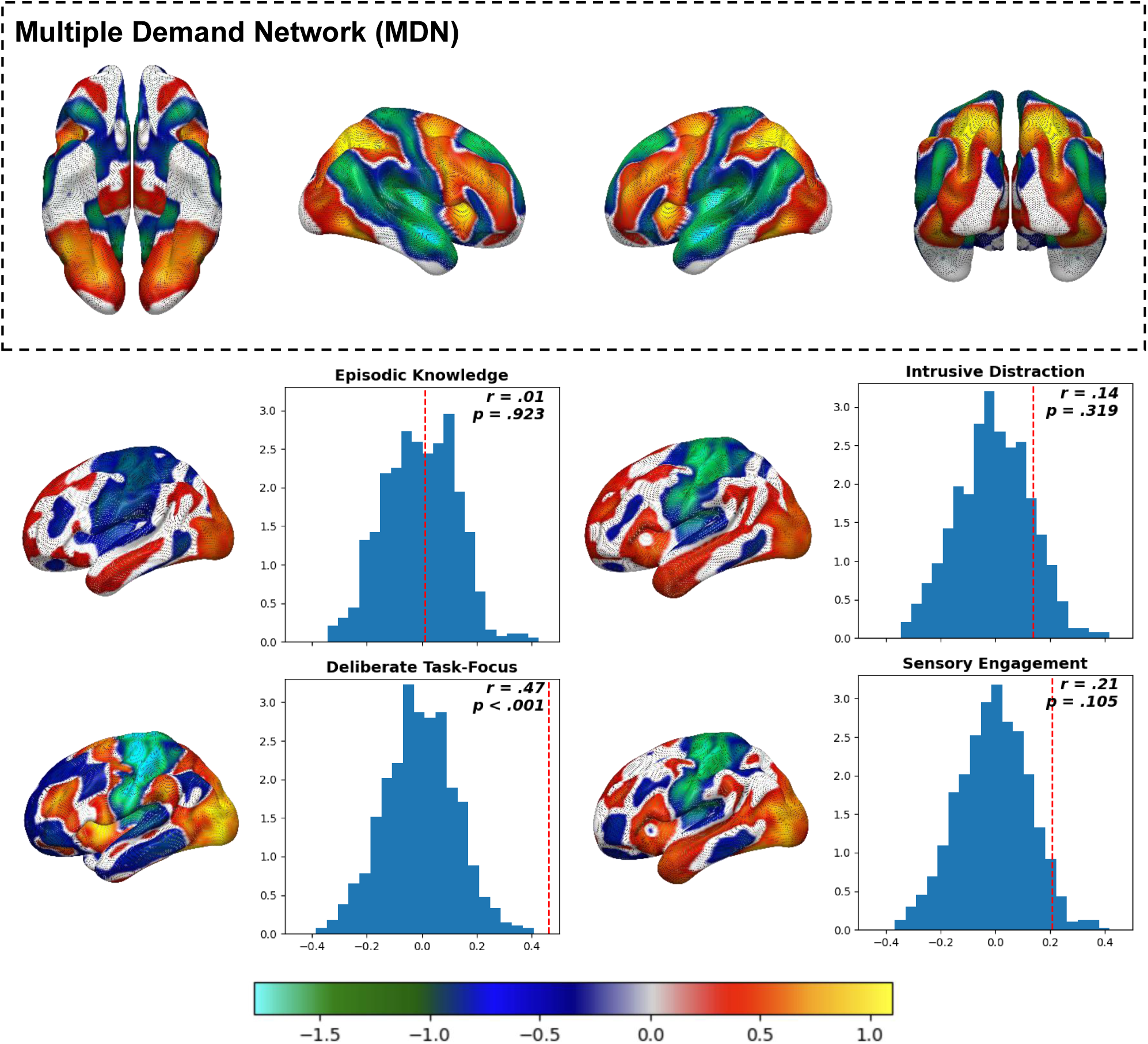
Regions related to stable *Deliberate Task-Focus* significantly overlap with the Multiple Demand Network (MDN). Participants’ reports of *Deliberate Task-Focus* were linked to stable task-related engagement and activation of brain regions implicated in executive functioning. As such, we sought to test the validity of our generated stability map by comparing it to the MDN, an established network with functional associations similar to *Deliberate Task-Focus*. Spin-testing found that the stability map for the component indeed significantly overlapped with the MDN, whereas the same effect was not present for any other thought pattern. This not only provides novel evidence supporting the association between the MDN and engagement with complex tasks but provides further nuance suggesting that as MDN activation grows, individual differences in *Deliberate Task-Focus* become more apparent.

Having established that task-level stability in reported thought covaried with both the content of experience and macroscale patterns of brain activation, we next examined whether these stability-linked patterns corresponded to any well-characterized functional systems. Because the *Deliberate Task-Focus* component was linked to both greater stability in probe responses and activation of control-related frontoparietal systems, we examined whether the resulting stability map for the component resembled the Multiple Demand Network (MDN) — a network consistently implicated in cognitive control and goal-directed task engagement (Duncan, 2010). Spin-testing (see Methods) revealed the stability map for *Deliberate Task-Focus* indeed overlapped with the MDN in a manner significantly greater than chance (𝑟 = .47, 𝑝 < .001; see Fig. 10). Similarity to the MDN was non-significant for all other thought patterns (*Episodic Knowledge*: 𝑟 = .01, 𝑝 = .923; *Intrusive Distraction*: 𝑟 = .14, 𝑝 = .319 ; *Sensory Engagement*: 𝑟 = .21, 𝑝 = .105), suggesting a selective link between stable reports of deliberate task-focused thought and the engagement of domain-general control systems.

### Results summary

We show that, across a diverse set of tasks, patterns of ongoing thought were not intrinsically stable properties of either individuals or task contexts alone. Instead, stability emerged from their interaction: the same experiential features were expressed consistently in some tasks but variably in others. Tasks that elicited higher levels of deliberate, task-focused thought showed greater within-subject stability, whereas tasks associated with intrusive distraction or strong sensory engagement tended to produce more variable reports of experience. Critically, task-level variation in the stability of deliberate task-focused thought was linked to patterns of whole-brain activation aligned with frontoparietal control systems. Stability maps derived from cortical gradients selectively overlapped with the multiple-demand network, a relationship not observed for other thought patterns. Together, these findings indicate that coordinated frontoparietal engagement is associated with the stabilisation of goal-directed cognitive states during task performance, linking large-scale brain organisation to the moment-to-moment consistency of ongoing experience.

## DISCUSSION

Our study examined how stability in ongoing thought emerges across different task contexts. There are four plausible ways that stable thought content could emerge: 1) some individuals could have very stable thoughts, 2) some task contexts could produce reliable thought patterns, 3) there could be some types of thought patterns that are intrinsically stable and 4) some types of thought patterns could be stable in some situations. Our findings provide limited support for accounts in which stable thought is attributable to individuals or tasks alone. Instead, stability emerged from their interaction: the same experiential features were expressed consistently in some tasks but variably in others. Tasks characterized by higher levels of deliberate, task-focused thought were associated with greater within-subject stability, whereas tasks associated with intrusive distraction or strong sensory engagement showed more variable reports. These findings indicate that stable modes of cognition are not fixed traits or task properties, but contextually supported states that depend jointly on the nature of the task, the individual, and the content of thought.

This pattern of results adds to an emerging body of research that highlights important contextually bound features of cognition^7,37^. In daily life, there are clear associations between the patterns of thought that individuals report and the activities they are performing. For example, ^1,2^ showed that activities such as work or homework can be reliably distinguished from activities like watching television or shopping based on the patterns of thought that individuals report. This context dependence of thoughts and actions can also be seen in how the weekly cycle shapes our thoughts, where individuals report more vocationally focused thoughts between 9 and 5 Monday to Friday than during the weekend^3^. Together, these findings show that patterns of thought are reliably shaped by situational context, supporting the view that cognition is dynamically organized around current activities rather than only reflecting fixed individual tendencies.

Lab studies have also provided important insights into how patterns of thought are reorganized in line with situational demands. For example, the relationship between cognitive traits such as fluid intelligence and ongoing thought can vary markedly across task contexts. Turnbull and colleagues showed that during a demanding 1-back task, individuals with higher fluid intelligence were better able to maintain task focus, whereas during a less demanding 0-back task, higher fluid intelligence was instead associated with greater engagement in task-unrelated, self-generated thought^38^. In the brain, this context-dependent regulation of thought has been linked to activity in the dorsolateral prefrontal cortex, a core region of MDN^4^. In a follow up study, Turnbull and colleagues established that the dorsolateral prefrontal cortex showed increased activity in the easier 0-back task when individuals engaged with self-generated thoughts about the future and the past, but that the same region showed increased activity when participants performed the more demanding 1 back task^20^. This indicates that frontoparietal control regions flexibly support different cognitive modes depending on situational demands.

Our findings provide converging evidence that regions of the MDN play a specific role in stabilizing deliberate, goal-directed modes of thought. Tasks in which deliberate task focus was reported most consistently were also those that showed the strongest engagement of regions within the MDN. This result suggests that one important function of the multiple demand system is to support the stability of thought, a possibility that logically flows from the argument that the MDN creates ‘cognitive episodes’^4^ which focus only on relevant information^39,40^, and must strengthen when tasks become more difficult^40^. Our data extend these accounts by demonstrating that multiple-demand engagement is associated not only with flexible control, but with the temporal stability of a cognitive state, supporting the endurance of deliberate task-focused thought across moments of ongoing performance. At the same time, these findings raise important questions about how stable cognitive states are best measured and how reliably they can be linked to underlying neural mechanisms.

### Questions to be addressed in future work

While this study advances understanding of how contextual factors shape cognitive states, several open questions remain. A key issue concerns the informativeness of introspective reports for characterizing cognitive function^41^. Although introspection offers unique access to internal experiences, its interpretive value depends on how reliably these self-reports can be situated within task contexts for which objective neural and behavioural measures are also recorded ^17^. Prior findings suggest that experience-sampling methods can capture stable, meaningful patterns of thought,^15,17,25,27,30,42–44^ while tools such as mDES have been shown to yield similar associations across countries^3,25^ and are sensitive to variation in age groups^45^. Encouragingly, the adaptability of mDES to smartphone-based administration presents a practical opportunity to broaden participation and mitigate the demographic biases that have long characterized psychological and neuroscientific research^32,46,47^.

Finally, we used a relatively novel analytic approach linking the reliability of thought patterns to brain activation patterns by regressing task-level variation in reliability onto coordinates in a lower-dimensional “brain-space” (see ^15,17,30^ for similar approaches). While this framework offers a promising bridge between behavioral and neuroimaging research, its validity and capacity to accurately predict brain states remain to be established. Replication and methodological refinement will be important for determining how robustly this approach captures whole-brain activation associated with stable cognitive states across situations.

## METHODS

### Participants

For our analyses we used a sample discussed in our previous publication ^17^. We recruited 194 participants from the undergraduate student population at Queen’s University, Canada, to complete a 14-task battery in our behavioral lab. We determined our sample size based on prior research investigating differences in ongoing thought across easy and hard task contexts. Our study received approval from the Queen’s University General Research Ethics Board, and we complied with all relevant ethical guidelines. All participants provided informed written consent and earned 2 course credits for their participation. Due to recording errors, we were unable to collect demographic data for four participants. This left us with a final sample of 190 participants: 164 identified as women, 24 as men, and two as non-binary or a similar gender identity. The average age of our participants was 18.56 years (SD = 1.09, range = 17–24 years). Each participant completed 38 mDES probes, contributing to a total of 7,220 observations across the study.

### Task-Battery

Participants attended a single 2-hour session in our behavioral laboratory to complete the 14-task battery. After providing written informed consent and filling out a brief demographic questionnaire, participants completed the battery alone in a room with a computer. To minimize distractions, we instructed them to avoid using any technological devices apart from the one provided for the study. The task battery (programmed using PsychoPy35) comprised 14 tasks, which we grouped into seven pairs based on task similarity. To improve clarity in our figures and writeup, we grouped our tasks into four categories based on their shared cognitive demands: 1) “Executive/Working Memory (WM)” tasks, which require relatively stronger concentration and recruit executive functioning processes, 2) “Visuomotor” tasks, involving decision-making based on immediate sensory information, 3) “Internally Oriented” tasks, where participants are prompted to introspect, often about information in autobiographical memory, and 4) “Naturalistic” tasks designed to emulate real-world activities. See Figure 1 for a summary of the set of tasks in each category.

Each task consisted of three blocks, except for the clip-viewing and Two-Back tasks, which included two blocks each, resulting in 38 task blocks in total. Task blocks were randomized within tasks (except for passive-viewing), each lasting ∼90s (with a ±15s jitter). Task pairs were presented consecutively, with task order within each pair randomized. Task pairs were also randomized based on a unique seed assigned to each participant. Written instructions preceded each task block, and after completing each block, participants answered mDES questions about their thoughts during the preceding block. The entire battery took approximately 1.5-2 hours to complete. Full code for the task battery is openly available on <u>our lab GitHub</u>.

## MULTI-DIMENSIONAL EXPERIENCE SAMPLING (MDES)

We measured participants’ ongoing thought using mDES probes administered at the end of every task block. Each mDES probe consisted of 16 items (see Table 3), presented in a randomized order. Participants rated each item on a continuous scale from one to ten by using the left and right arrow keys to adjust a marker on the screen and pressed enter to submit each response. To avoid bias, the marker’s starting position was randomized for each item. Participants completed 38 probes over the course of the battery.

**Table 3.**
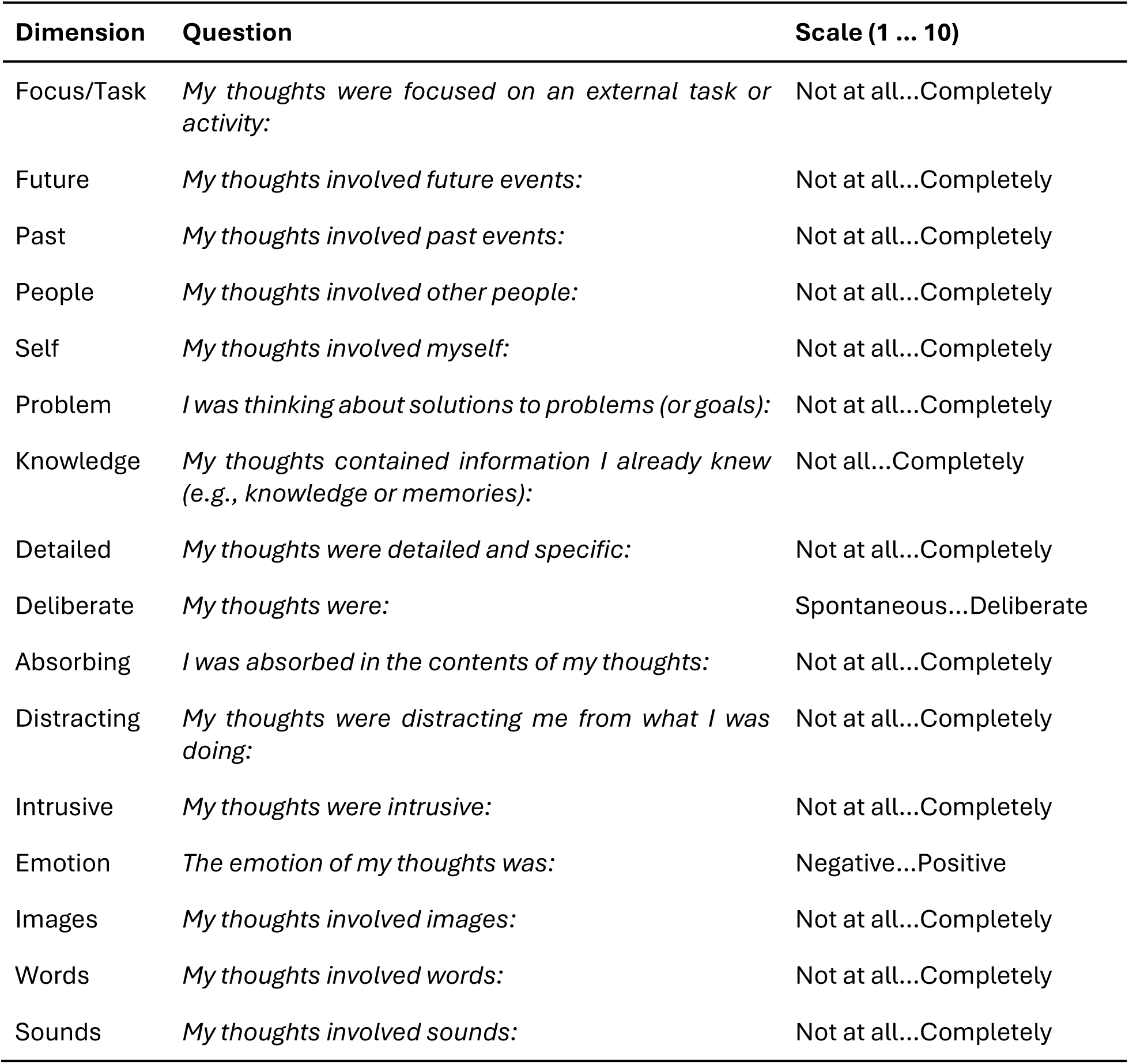
Multidimensional Experience Sampling (mDES) Items.

## EXECUTIVE/WORKING MEMORY TASKS

### Math Task

Adapted from ^18^ who used it as a localizer task for the Multiple Demand Network (and by deactivation, the DMN). Participants viewed addition expressions and were instructed to use a keypress to select the correct sum from two options. Each expression appeared for 1.45s, followed by the two sums for 1.45s. In the ’easy’ condition, expressions involved one-digit numbers, while the ’hard’ condition included one two-digit number (up to 19). Each block comprised 28 trials. If no response was made within 5s, the trial was counted as incorrect.

### 2-Back Tasks

We designed this task to mimic the 2-back portion of working memory/cognitive control task from the HCP ^19,48^. Participants pressed a key to indicate whether a presented stimulus matched the one shown two trials prior. We used two of the 4 types of stimuli from the HCP: faces (“2B Faces”) and scenes/places (“2B Scenes”) ^19^. Participants were told to “track whether the image presented at any point is the same as the one two images before it” and to use the left and right arrow keys to indicate the image as matching or not, respectively. Each 2-back task had two blocks, each containing 35 trials, including five ’target’ trials (where a stimulus repeated after two trials) and five ’target-lure’ trials (where a stimulus repeated after either 1 or 3 trials). Each trial lasted 2s, with a 500ms inter-stimulus interval.

### 0/1-Back Task (0-Back: Visuomotor; 1-Back: Executive/WM)

We adapted this task from ^20^, who used it to compare neural activation and reported thought between task conditions requiring different levels of focus. Participants viewed non-target trials where two different shapes (e.g., square and circle) appear on either side of a central black line for 0.5-1.5 seconds. After 2-8 non-target trials, a target trial would present a smaller shape over the line for 3.5-5 seconds (or until input). In the 0-back condition, participants indicated with a keypress which of two shapes presented on the left and right matched the central shape. In the 1-back condition, participants indicated the matching shape from the previous non-target trial. A fixation cross appeared for 1-2.5s between trials.

Since we were more interested in characterizing the unique states linked to each condition, we administered the 0- and 1-back conditions in separate blocks rather than presenting them in an alternating or counterbalanced fashion. As each condition relies on different cognitive demands we grouped the isolated conditions into separate task categories. Namely, we categorized the 1-back as “Executive/WM” because it relies on maintaining information from non-target trials in working memory, whereas the more immediate perceptual decision-making during the 0-back better fits into the “Visuomotor” category.

## VISUOMOTOR TASKS

### 0-Back Task

We describe this task above together with its associated 1-back condition.

### Finger-Tapping Task

We modeled this task on the finger-tapping portion of the motor task used in the Human Connectome Project (HCP) ^19^. Participants were told to press the spacebar when a black square appeared on the screen, with a fixation cross shown between presentations. Participants were told to use their right or left hands across different blocks. Squares remained visible for 2s, and missing a response within this time was recorded as a failure. The delay between squares was jittered, ranging from 2-5s.

### Go/No-Go Task

Participants responded with a button press for ‘Go’ trials or withheld their response for ‘No-Go’ trials. We adapted this task from ^21^, who sought to compare neural activation when participants withheld responses according to perceptual versus semantic criteria at varying levels of difficulty. We only used the perceptual condition, where participants chose responses according to the degree of slant in a rhombus. Each trial presented a scrambled word framed in a rhombus outline for 0.75–1.25s. We instructed participants that a slight slant to the rhombus indicated a ‘Go’ trial, while a larger slant signaled a ‘No-Go’ trial. Our study used only the ‘hard’ condition from the original task, where slants were harder to distinguish. A fixation cross appeared between trials for 0.5–1s. Each block included 46–54 stimuli with 80% ‘Go’ and 20% ‘No-Go’ trials.

## INTERNALLY ORIENTED TASKS

### Trait-appraisal/“Reference” Tasks

In these tasks, which we adapted from ^22^, we asked participants to evaluate whether an adjective “applies to” themselves (“Self-reference” aka “the You Task”) or “your best friend” (“Social reference” aka “the Friend Task”). In the original study, the authors examined whether DMN activation during this task predicted *any* self-/other-relevant thought when off-task. As such, participants in the original study made judgments about some familiar person rather than a close friend, specifically. Adjectives appeared one at a time in the center of the screen and were either positive (e.g., “enthusiastic”) or negative (e.g., “insecure”). Participants used the left and right arrow keys to indicate whether they associated the adjective with the referent or not. For the “Friend” condition, participants were asked to consistently think of the same friend throughout the task. Each block included a unique, counterbalanced set of adjectives across tasks.

### Memory Task

We adapted this task from ^23^, who aimed to distinguish the default-mode activation associated with reading and autobiographical recall. In their study, participants were either asked to read sentences presented one word at a time, or to recall a personal experience prompted by a one-word cue. We divided the recall and reading conditions into separate tasks in our battery. In each block, participants viewed a cue-word (e.g., ’Airport’) and were then asked to recall a memory related to it, pressing ’Enter’ when they had one. Afterward, a 4-5s fixation cross appeared, followed by instructions: “Now we would like you to think about this event. Please press Enter when you are ready to begin.” A 20s fixation cross was then shown, followed by an mDES probe.

## NATURALISTIC TASKS

### Movie Viewing

Participants watched clips (3:46 in duration) from either *Inception* (‘Sci-Fi Clip’) or *Welcome to Bridgeville* (‘Documentary Clip’) taken from the set of clips used in movie-viewing scans from the HCP ^24^. We balanced the clip volumes to minimize noise spikes. For each clip, we probed subjects twice: once between the first and last 15s of the clip, and a second time at the clip’s conclusion.

### Reading Task

We adapted this task from the reading task from ^23^ (see Memory Task). During the reading task, participants read 15-word sentences, with words presented one at a time. After a 1-3s fixation cross, each word appeared for 600ms.

### Task Brain Maps

The spatial maps summarizing brain activity for each of the 14 tasks were drawn from prior fMRI studies (See table on the left side of Figure 1 for the source publications for each Task Map). These unthresholded, group-averaged, *z*-stat contrast maps compared task conditions to baseline, representing the group-averaged BOLD signal for each task. To account for differing coverage across maps, we applied a binarized mask that removed regions with missing data and used it for all maps before projection into gradient space. All brain maps, including the gradient maps, are available in the <u>StateSpace GitHub repository</u>.

### Statistical Analysis

#### DECOMPOSITION OF EXPERIENCE SAMPLING PROBES

Per previous work (see ^32^ for a review; ^1,2,15,25,29^, we obtained the dimensions of the “thought-space” by reducing subjects’ mDES probe responses using Principal Components Analysis (PCA). The output components of the PCA represent patterns of covariance among mDES items that capture large chunks of variance in reported thought. Each of these “thought patterns” forms a dimension of the subsequent thought space. To determine the number of components to extract we used Parallel Analysis^49^ and examined the slope of the Scree plot (See Fig. 2). To maximize interpretability, we applied Varimax rotation to the set of components and labelled the resulting dimensions based on their loading structure. We then estimated Regression component scores for each mDES probe on each component.

#### DIMENSION REDUCTION OF BRAIN MAPS

To represent the whole-brain activation associated with each task in a smaller number of dimensions, we projected each map onto a set of five connectivity gradients generated by ^33^ using the averaged functional connectivity matrix of the resting state data collected during the Human Connectome Project (See the study’s <u>collection on Neurovault</u> for cortical and subcortical maps of each gradient; See Fig. 4). Margulies et al. (2016) generated these dimensions using diffusion embedding, a dimensionality reduction technique that identifies large-scale patterns in whole-brain connectivity (termed ‘gradients’), which it returns in descending order of variance accounted for. Plotted on these gradients, regions with similar connectivity profiles tend to fall closer together, while those with more distinct profiles fall farther apart. The first five gradients explain about 60% of connectivity variance, and prior research suggests that the first three relate to important characteristics of each brain region’s functional profile. More specifically, the first gradient (unimodal—transmodal) differentiates sensorimotor cortex from association cortex, the second (motor—visual) distinguishes motor from visual cortex, the third (DMN—control) separates the default mode network from areas linked to executive control (e.g., the frontoparietal control network), the fourth (DAN—sensory attention) differentiates the dorsal attention network from the ventral attention network and visual cortex, and the fifth (sensory motor—Limbic/VAN) distinguishes visual cortex from the ventral attention network (see Fig. 4).

We estimated ‘gradient scores’ for each of the 14 task maps on each of the 5 connectivity gradients, by calculating pairwise spatial correlations (Spearman rank) between each masked task brain map and each gradient. This process yielded five correlation values for each task map, with each coefficient representing its position along each gradient dimension (See Fig. 5). For example, since the first gradient separates sensorimotor on the negative end from association cortex on the positive end, a stronger positive correlation with gradient 1 would indicate that a brain map comprises more activation of association cortex than sensorimotor cortex, while a stronger negative correlation would indicate the opposite. These correlation values thus serve as ‘coordinates’ in the 5-dimensional space. The code for this analysis is available on the <u>StateSpace Github</u>.

#### QUANTIFYING STABILITY IN REPORTED THOUGHT

We used the Intraclass Correlation Coefficient (ICC) to examine how participants’ descriptions of their thoughts varied in stability across probes. The ICC represents the proportion of the total variability in a measure captured by variability between-subjects ^35,50^. A measure thus yields a higher ICC the more it: 1) consistently characterizes individual levels of a grouping variable and 2) strongly differentiates unique levels ^50,51^. In other words, measures with a higher ICC more effectively distinguish between members of a group. We visualize this ratio of between- to within-unit variability using ^36^’s *theoretical variation field map*. The field map plots between-subject variation on the y-axis and within-subject variation on the x-axis. Capturing the above idea, points yield higher ICC values as they approach the top-left corner of the map, where the ratio of between- to within-subject variability is large and measurements are the most reliable (See Fig. 7) ^36^. For example, a task with both low between-subject variability and low within-subject variability would fall in the bottom left corner of the map. In this case, subjects score consistently on the task, but in a highly similar manner. This region thus describes individuals with only moderate reliability, since subjects provide reliable responses to the task but are difficult to tell apart from each other. This could be the case for situations that exert a strong influence over internal experience, constraining individuals to a smaller range of responses.

We examined reliability among subjects’ reported thoughts at three levels: we first examined the repeatability of subjects’ scores across tasks (subject-level ICCs), then the repeatability of probe scores within tasks across subjects (task-level ICCs), and finally the repeatability of subjects’ scores across probes within each task (subject x task-level). In all cases we used the bootstrapICC function in R 4.2.2 ^52^ to generate bootstrapped estimates for the ICC for each component with 95% confidence intervals, which uses the lme4 package ^53^ for mixed-effects modelling and the boot ^54^, and lmeresampler packages ^55^ for bootstrapped resampling.

##### Trait-Like Reliability: Stability across tasks within subjects

To examine how the reliability of individuals’ probe responses varied *independent* of the task at hand, we computed ICCs using a 1-way mixed effects model for each rotated component with each model including a random-intercept for subject, i.e.:

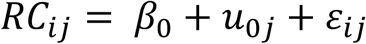

where 𝑅𝐶_𝑖𝑗_is the score on a given rotated component for probe 𝑖, subject *j*. 𝛽_0_ is the grand mean of the component scores, and 𝑢_0𝑗_ ∼ 𝑁(0, 𝜎_u_^2^) is the random intercept for subject 𝑗.

Written in R formula format:

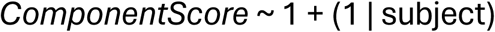

This ICC can also be thought of as the average correlation between any two probes randomly sampled within a subject across the set of tasks. According to this measure, stability in a given thought pattern is disconnected from task-context and more a result of intrinsic characteristics of the participant. As such, we refer to this as a ‘trait-like’ ICC.

##### State-Like reliability: reliability across subjects within tasks

Complement to estimating variance that can be attributed to differences between subjects, we also calculated variance that can be attributed to differences between task conditions. In this case, a high ICC indicates that tasks elicit reliable probe responses that differ between tasks. For example, a high task-level ICC for, say, task-focus, suggests that each task stably yields its own unique level of reported task-focus. We also computed these ICC estimates using a one-way mixed effects model for each rotated component but this time we instead included a random intercept for task-condition, i.e.:

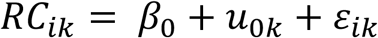

where 𝑅𝐶_𝑖𝑘_ is the score on a given rotated component for probe 𝑖, task 𝑘, 𝛽_0_ is the grand mean of the component scores, and 𝑢_0𝑘_ ∼ 𝑁(0, 𝜎_u_^2^) is the random intercept for task 𝑘.

Written in R formula format:

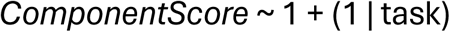

As this ICC grows, subjects respond more similarly within each task. As such, the ICC can also be conceptualized as the average correlation between two probes randomly sampled from a given task condition. Since this measure captures the change in component-score attributed to participants inhabiting a particular task-state, we refer to this as a ‘state-like’ ICC.

State x Trait reliability: within-subject reliability per task

Finally, in addition to examining how reliability varied depending on the subject and task independently, we also analyzed their interaction (i.e., task-*dependent* within-subject reliability). For this we generated overall ICCs using a one-way cross-factored random effects model for each rotated component with nested random intercepts for each subject-task pairing, e.g.:

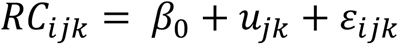

Where 𝑅𝐶_𝑖𝑗𝑘_ is the rotated component score for probe 𝑖, during task 𝑘, for subject 𝑗, and 𝑢_𝑗𝑘_ ∼ 𝑁(0, 𝜎_*u*_^2^) is the random intercept for the combination of task 𝑘 with subject 𝑗. In R, we input this as:

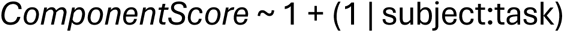

Unlike the above ICCs, which describe reliability within-subjects averaged across tasks or within-tasks averaged across subjects, this ICC measures how subjects respond more reliably *depending* on the task at hand. In other words, how reliably can we identify subjects based on their responses if we take into account what they are doing. This ICC value grows as subjects’ responses become more stable and unique within each task. It can thus also be conceptualized as the average correlation between probes randomly sampled from a participant within a given task.

To elaborate on this interaction further, we used univariate models to generate subject-level ICCs for each task, representing how within-subject stability and between-subject differences changed for each condition. With this measure, a higher subject-level ICC suggests subjects consistently think uniquely from each other during a given task. A lower subject-level ICC, on the other hand, suggests that subjects’ responses are *not* consistently different from each other. Note that a low ICC could either reflect that variability is either too low (i.e., subjects are highly similar) or too high (i.e., measurements are too unreliable) for individual differences to be evident.

#### THOUGHT STABILITY AND COMPONENT-SCORE

To examine whether task-level stability in probe responses varied systematically with thought content, we tested the relationship between stability on each component across tasks and component score. For each thought pattern, we fit linear models regressing the ICCs for each task on task-averaged component scores. In each model we used Fisher *z*-transformed ICCs so that the variate would be normally distributed^56^. Because these analyses only used 14 task-level observations, we used bootstrap resampling to assess the robustness of each relationship rather than relying on statistical significance. In each of 1,000 resamples, participants were resampled with replacement, task-level ICCs and component averages were recomputed, and the regression model was refit. From the resulting distributions, we estimated the median regression coefficient and percentile-based 95% confidence intervals. We also estimated the directional consistency of each effect by computing the percentage of coefficients across resamples with a positive versus negative slope.

#### THOUGHT STABILITY AND WHOLE-BRAIN ACTIVATION

In addition to relating task-evoked patterns of thought to the stability of probe responses, we examined how task-level stability was associated with whole-brain activation. To account for shared variance between gradients, we fit bootstrapped multiple regression models (𝐵 = 1000) for each thought component, using Fisher’s z-transformed ICCs as the outcome and task coordinates on all five cortical gradients as predictors. As in the preceding analyses, resampling was performed at the participant level with replacement. Because our interest lay in how this relationship varied relative to the average across the task battery, task-level gradient coordinates were z-scored prior to model fitting. From the bootstrap distributions, we estimated median regression coefficients for each gradient–component pairing and percentile-based CIs, corrected for multiple comparisons using a Bonferroni adjustment. Here too, directional consistency was quantified as the proportion of bootstrap resamples yielding positive versus negative slopes.

Under this framework, a stronger association between task-level stability for a given thought component and position along a gradient indicates that tasks occupying a particular region of gradient space are associated with more stable, person-specific expression of that mode of thought. For example, a positive relationship between the stability of responses to the *Detail* item and the unimodal-transmodal gradient would suggest that tasks engaging association cortex more strongly elicit more internally consistent, trait-like reports of detailed thought.

#### GENERATING ‘STABILITY MAPS’

To better visualize the set of relationships between task-dependent stability and scores on each of the 5 principal whole brain gradients, we multiplied each gradient by its median bootstrapped regression coefficient predicting stability on a given component and then averaged across the 5 re-weighted gradients to generate dot-product maps. In equation form:

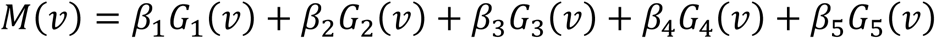

where 𝑀(𝑣) is the voxel weighting for a given component’s stability map at voxel 𝑣, 𝛽_𝑘_is the regression coefficient relating the coordinates for gradient 𝑘 to the component’s ICC per task and 𝐺_𝑘_(𝑣) is the *z*-scored weighting for gradient 𝑘 at voxel 𝑣. We generated these maps in Python 3.14.1 using the pandas^57,58^ and NumPy^59^ libraries.

These stability maps gave greater weight to vertices more strongly associated with stability on a component, effectively producing a beta-map for stability on each thought pattern (See ^17^ for an analogous approach).

#### SPIN-TESTING

To test the similarity of each stability map with the multiple-demand network (MDN), we conducted permutation testing by generating 1000 random spin permutations ^60^ of each stability map on a sphere and generated a null distribution by computing the Spearman correlations of each of these null permutations with the MDN. We then obtained the p-value for the true correlation estimate on the generated spin distribution. We represented the MDN using a surface map template of the core and extended multiple demand network generated with data from the Human Connectome Project (HCP)^24,61^. We conducted this analysis in Python 3.14.1 using the neuromaps package^62^.

## Supporting information

Supplementary Results

