## Supplementary Results for "The stability of thought: using experience sampling and brain imaging to determine the contextually bound nature of human cognition"

### Trait- versus state-like stability across different thought patterns

To examine how different aspects of thought were differently driven by individual- versus task-level differences, we computed marginal ICCs for each thought-pattern nesting participants' probes either within subject or within task. Higher within-subject ICCs indicate that a given thought-pattern scored relatively consistently within subjects and uniquely between subjects, while higher within-task ICCs suggest that a given thought-pattern scored more uniformly within a task across subjects and uniquely between tasks. In other words, components with higher subject-level ICCs more effectively distinguish individuals, while components with higher task-level ICCs more effectively distinguish task contexts.

| **Thought-Pattern** | **Subject ICC** | **Task ICC** | **ΔICC$_{(\text{Subject – Task})}$** |
|---|---|---|---|
| *Episodic Knowledge* | .17 [.13, .20] | .36 [.19, .55] | -0.19 |
| *Intrusive Distraction* | .37 [.31, .42] | .04 [.00, .07] | 0.33 |
| *Deliberate Task-Focus* | .33 [.28, .38] | .13 [.04, .20] | 0.20 |
| *Sensory Engagement* | .28 [.23, .33] | .20 [.07, .31] | 0.08 |

**Table S1*.*** ICCs for each thought-pattern nested within-subject, within-task, and the difference in ICC values for each component. ICCs are presented with bootstrapped 95% confidence intervals.

Examination of the within-subject and -task ICCs for each thought pattern revealed that the variance accounted for by each factor differed depending on the thought-content in question (see Table S1). Most notably, the first two components, *Episodic Knowledge* and *Intrusive Distraction*, demonstrated inverse compositions of subject- and task-level variation. Specifically, *Intrusive Distraction* exhibited the highest ICC by subject ($ICC_{subject} = 0.37$), but the lowest ICC by task ($ICC_{task} = .04$), whereas *Episodic Knowledge* associated with the opposite pattern ($ICC_{subject} = .17$; $ICC_{task} = .36$). Put differently, *Intrusive Distraction* scores were on average more correlated when sampled from within a given subject than sampled within a given task, whereas the opposite was the case for *Episodic Knowledge*. *Sensory Engagement* ($ICC_{subject} = .28$; $ICC_{task} = .20$) and

*Deliberate Task-Focus* ($ICC_{subject} = .33$; $ICC_{task} = .13$) also showed differences in trait- vs state-operation, varying more due to subject differences than task differences, though to a smaller extent than *Intrusive Distraction*.

In sum, although dependencies on strictly task or subject were modest, comparison of within-task vs within-subject ICCs across components revealed that certain thought patterns seemed to fluctuate more as a result of trait-like factors independent of surrounding context (particularly *Intrusive Distraction*), while other characteristics of thought (e.g., temporal-orientation, other-orientation, recruitment of previous knowledge, etc.) appeared more task-*dependent*, varying more due to the task at hand.

**Bootstrapped regression of ICC against gradient coordinates**

| Thought Pattern | Median *b* | $CI_{adj.}$ | Directional Consistency |
|---|---|---|---|
| **Episodic Knowledge** | | | |
| *Uni.-Hetero.* | 0.04 | [-0.03, 0.09] | 97.2% |
| *Mot.-Vis.* | 0.07 | [-0.09, 0.23] | 89.7% |
| *DMN-Cont.* | -0.01 | [-0.09, 0.06] | 63.2% |
| *VAN-S.Att.* | 0.05 | [-0.05, 0.16] | 93.5% |
| *S.Mot.-Lim.VAN* | -0.01 | [-0.16, 0.16] | 60.2% |
| **Intrusive Distraction** | | | |
| *Uni.-Hetero.* | 0.07 | [0.01, 0.14] | 99.9% |
| *Mot.-Vis.* | 0.11 | [-0.02, 0.25] | 99.5% |
| *DMN-Cont.* | 0.04 | [-0.05, 0.12] | 91.0% |
| *VAN-S.Att.* | 0.08 | [-0.01, 0.17] | 99.4% |
| *S.Mot.-Lim.VAN* | 0.04 | [-0.08, 0.17] | 85.4% |
| **Deliberate Task-Focus** | | | |
| *Uni.-Hetero.* | 0.03 | [-0.03, 0.09] | 89.8% |
| *Mot.-Vis.* | 0.19 | [0.03, 0.34] | 100.0% |
| *DMN-Cont.* | 0.11 | [0.02, 0.19] | 100.0% |
| *VAN-S.Att.* | 0.08 | [-0.01, 0.18] | 98.9% |
| *S.Mot.-Lim.VAN* | 0.09 | [-0.04, 0.23] | 97.7% |
| **Sensory Engagement** | | | |
| *Uni.-Hetero.* | 0.04 | [-0.01, 0.09] | 98.1% |
| *Mot.-Vis.* | 0.14 | [-0.02, 0.30] | 99.6% |
| *DMN-Cont.* | 0.03 | [-0.04, 0.12] | 86.9% |
| *VAN-S.Att.* | 0.06 | [-0.02, 0.15] | 98.7% |
| *S.Mot.-Lim.VAN* | 0.07 | [-0.07, 0.22] | 91.1% |

**Table S2. Full bootstrapped regression results regressing task-level gradient coordinates against task-level ICC values for each thought pattern.** Median regression coefficients for each component-gradient combination with Bonferroni-corrected percentile-based bootstrapped *CI*s ($CI_{adj.} = 99.75\%$). The 'directional consistency' of each effect refers to the percentage of resampled coefficients with the same sign as the median coefficient.
